# Understanding the mechanism of the SARS CoV-2 coinfection with other respiratory viruses

**DOI:** 10.1101/2022.12.15.520197

**Authors:** Kalaiarasan Ponnusamy, RadhaKrishnan Venkatraman Srinivasan, Robin Marwal, Meena Datta, Mahesh Shankar Dhar, Urmila Chaudhary, Uma Sharma, Swati Kumari, Kalpana Sharma, Hema Gogia, Preeti Madan, Sandhya Kabra, Partha Rakshit

## Abstract

Coinfections have a potential role in increased morbidity and mortality rates during pandemics. Our investigation is aimed at evaluating the viral coinfection prevalence in COVID-19 patients. Rapid diagnostic tests are tools with a paramount impact both on improving patient care. Particularly in the case of respiratory infections, it is of great importance to quickly confirm/exclude the involvement of pathogens. The COVID-19 pandemic has been associated with changes in respiratory virus infections worldwide, which have differed between virus types. In this paper, we systematically searched the percentage of coinfection of various respiratory viruses in COVID-19-positive samples. We included patients of all ages, in all settings. The main outcome was the proportion of patients with viral coinfection. By describing the differences in changes between viral species across different geographies over the course of the COVID-19 pandemic, we may better understand the complex factors involved in the community cocirculation of respiratory viruses.

## Introduction

Accumulating evidence reveals that the high prevalence of co-infection associated with respiratory viral infections is the potential to worsen the clinical outcome of the disease [1]. However, few data available to date on co-infection have represented the minimum of the global population and examined the small number of viruses [2–4]. Also, the overlapping symptoms make the diagnosis difficult and may also lead to false diagnosis and missing coinfection cases, which may lead to a rise in co-epidemics and overburdening of the health care system [1, 5]. Molecular tools are the gold standard methods used to diagnose viral respiratory pathogens [6]. To date, Polymerase Chain Reaction (PCR) assays are the most used diagnostic tool to identify the viral respiratory pathogen [7]. These assays are rapid, relatively inexpensive, sensitive, and can be combined to identify multiple pathogens in a single test. However, drawbacks associated with this technique are limited to the number of targets included in the assay, therefore new or emerging pathogens will evade detection by PCR. Furthermore, the sensitivity of the assay is reduced due to mutation in the targeted region that leads to fail in detection or false negative results [6]. Introducing next-generation sequencing (NGS) could resolve these issues and revolutionize respiratory viral diagnostics and simultaneously analyze their genetic sequence. In contrast to other methods used for viral detection, NGS offers numerous advantages such as understanding the viral evolution, determining the source of infection and route of transmission identifying and characterizing the co-infections, and screening targets for possible therapeutics [6, 8]. Metagenomic sequencing using NGS technology is capable of identifying a variety of pathogens with the limitation of a large number of non-specific sequencings [9]. Alternatively, novel panels are available that use different oligo probes to enrich sequence targets of multiple respiratory DNA and RNA viruses that reduce non-specific reads in NGS data and provide high performance of targeted sequencing-based pathogen identification. In this study, we have analyzed a few positive and negative samples of Severe Acute Respiratory Syndrome Coronavirus 2 (SARS-CoV-2) for the identification of coinfection of other respiratory viruses using the Respiratory Virus Oligo Panel (RVOP) of Illumina.

## Materials and Methods

### Samples

In this study VTM samples collected from different health facilities of Delhi and around areas of Delhi were tested for SARS CoV-2 using Roche COBAS 6800, a fully automated real time PCR machine. Positive samples of SARS CoV-2 were further subjected to sequencing using Illumina sequencer Nextseq550. RNA of positive samples was extracted using an automated Roche MagNA Pure 96 system.

### Next-generation sequencing

RNA extracts were used as input for library preparation by using RNA Prep with Enrichment kit (Cat No: 20040537, Illumina). The following steps were involved in the process: first and second-strand cDNA synthesis, double-stranded cDNA tagmentation by bead-linked transposomes (BLT) and purification. The tagmented fragments were amplified to add index adaptor sequences using single-use Illumina^®^ DNA/RNA UD Indexes (Cat No: 20027213, Illumina). After Clean-up, libraries were quantified using Invitrogen Qubit dsDNA broad range Assay Kit and normalized according to the manufacturer’s instructions. Individual libraries were then combined in 3-plex reactions for probe hybridization using oligos from the respiratory panel (Cat No: 20044311, Illumina). The Respiratory Virus Oligo Panel (RVOP) of Illumina was used for probe hybridization which features ~7800 probes designed to detect respiratory viruses, recent flu strains, and SARS-CoV-2, as well as human probes to act as positive controls in every reaction. Hybridized probes were then captured, washed, and amplified. Library quantities and fragment size were determined using a Qubit 1×dsDNA HS assay and Agilent HS Tapestation respectively and sequenced using mid-output sequencing kit 2 × 75-bp runs on the Illumina Nextseq550 sequencer.

### Bioinformatic Analysis

After the sequencing using the NGS platform, bcl2fastq software is used to demultiplexes data and convert BCL files to Fastq files. The data analysis is carried out using the DRAGEN Pathogen Detection pipeline. Here we used the combined reference of humans and viruses to analyze pathogen data and create consensus FASTAs.

### Results and Discussion

The NGS analysis showed coinfection of SARS-CoV-2 with other viruses. Sequencing the SARS-CoV-2 genome to identify novel variants that can provide actionable information to prevent or mitigate emerging viral threats, predict transmission/resurgence of regional outbreaks, and also to test for co-circulating respiratory viruses that might be independent factors contributing to the global disease burden.

The aim of the study was to find and illustrate the correlation between SARS-CoV2 and coinfecting organisms and their disease manifestation. For the identification of coinfections, we used the NGS approach of target amplification and probe hybridization (RVOP, Illumina). VTM samples collected from different health facilities of Delhi and around areas of Delhi were tested for SARS CoV-2. Total 288 positive samples were further subjected to sequencing using RVOP panel in Illumina sequencer Nextseq550. The NGS analysis showed the coinfection of SARS-CoV-2 with other viruses. Out of 288 samples analysed, 17 samples (6%) were detected with 6 human respiratory viruses other than SARS-CoV2 (Figure 1). The human respiratory viruses such as Influenza A virus (A/Texas/50/2012(H3N2)), Influenza A virus (A/New York /392 /2004 (H3N2)), Human coronavirus 229E, Human rhinovirus A89, Human coronavirus HKU1 and SARS coronavirus Tor2 were mainly present as coinfections with SARS-CoV2 (Figure 2). Influenza A virus (A/Texas/50/2012(H3N2)) were present in 5 of 17 samples whereas Influenza A virus (A/New York /392 /2004 (H3N2)) present in 7 of 17 samples of SARS-CoV2 patients. Human coronavirus 229E and Human coronavirus HKU1 detected in 3 of 17 samples, whereas SARS coronavirus Tor2 in 2 samples and Human rhinovirus A89 in 1 sample were identified as coinfecting agents (Figure 2). Sequencing SARS-CoV-2 genome to identify novel variants that can provide actionable information to prevent or mitigate emerging viral threats, and predict transmission/resurgence of regional outbreaks and also to test for co-circulating respiratory viruses that might be independent factors contributing to the global disease burden.

**Figure 1.**
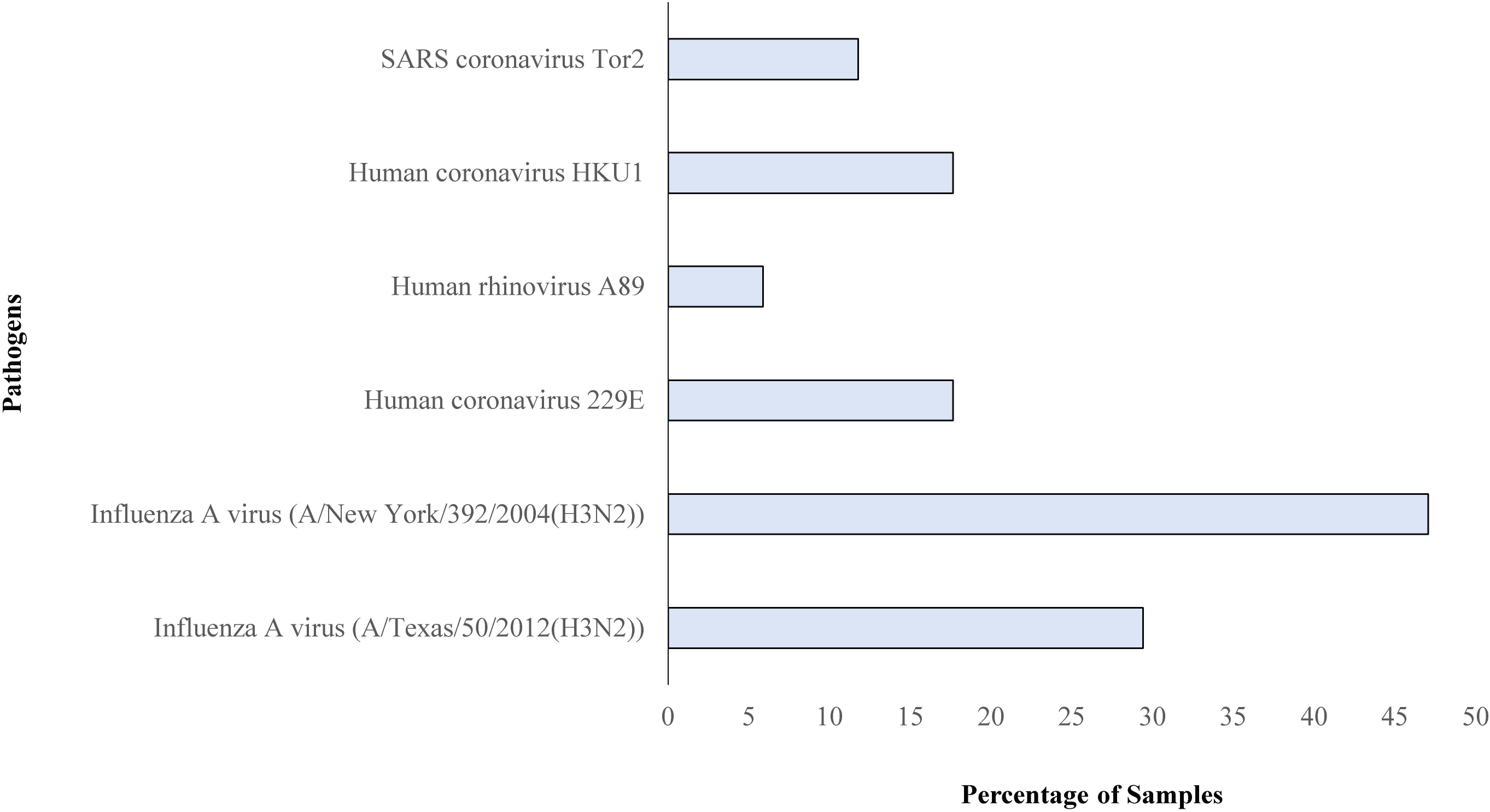
The RVOP panel virus coinfection with SARS-CoV-2

**Figure 2.**
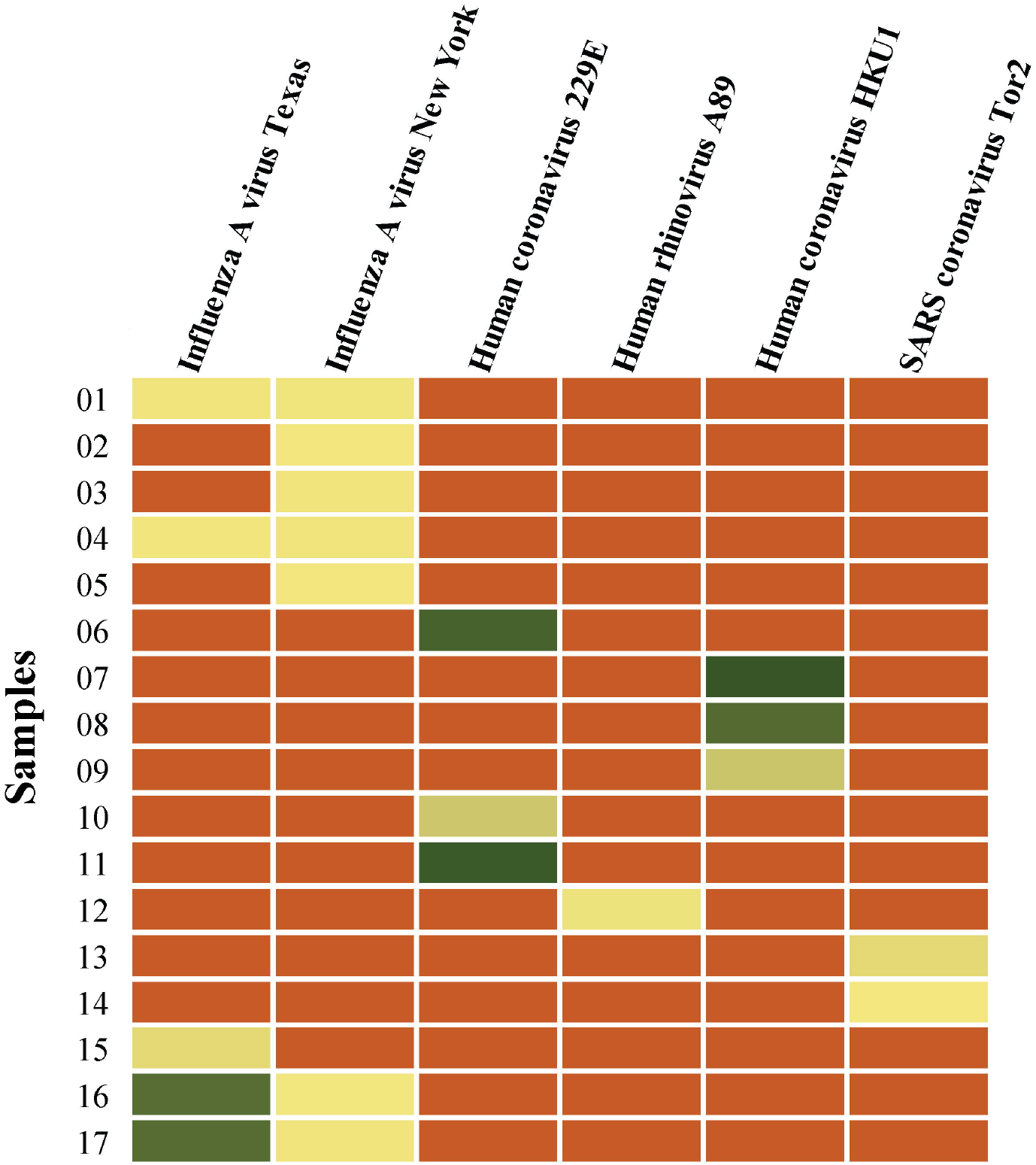
Coinfection of panel virus with SARS-CoV-2 positive samples

The present study utilized NGS approach of target amplification and probe hybridisation (RVOP, Illumina) to identify the co-infecting species present within the upper respiratory tract of SARS-CoV2 patients from Delhi and around areas. This method can achieve viral detection sensitivity comparable with molecular assay and also obtain partial to complete genome sequences for 41 respiratory viruses including their subtypes to allow accurate genotyping and variant analysis [10]. We used nasopharyngeal and oropharyngeal swabs to identify the respiratory tract microbiota that cooccurred with SARS-CoV2 patients. In this study, out of 288 samples analysed, 17 samples were detected with 6 human respiratory viruses other than SARS-CoV2. Mainly, Influenza A virus (H3N2) were coinfected with SARS-CoV2 in compare to other respiratory viruses. Both Influenza A/New York/392/2004 (H3N2) and Influenza A/Texas/50/2012 (H3N2) variant were prevalent in most positive samples of around 13 of 17 samples (75%). Coinfection of SARS-CoV2 with Influenza virus is very common due to seasonality in which known higher circulation of Influenza virus happened and also because of lower usage of Influenza vaccination in the specific area [11]. Few Human coronavirus 229E were present in this study. Human coronavirus 229E is known to be an opportunistic pathogen which cause life threatening infections in immunocompromised individuals [12]. Human rhinovirus found in lesser number of samples. It has been suggested that Human rhinovirus may reduce the infections of SARS-CoV2 if coinfection occurs simultaneously [13]. The findings of this study have tried to correlate the SARS-CoV2 infections with other respiratory viruses for disease monitoring. In this study we used the Respiratory Virus Oligo Panel (RVOP) of Illumina. The method is expected to detect many commonly known respiratory viruses. The high capture efficiency with high sensitivity from this method makes it a powerful tool for the discovery of respiratory virome in case of co-infections. Apart from RVOP, other sequencing panels are also available such as the Respiratory pathogen ID/AMR panel that helps to detect 180+ bacteria, 40+ viruses, 50+ fungi, and 1200+ AMR alleles respectively at a single time along with the genome sequences of viruses and other pathogens. The newly available Virus Surveillance Panel offers the detection of 66 pathogenic viruses that are of high risk to public health including SARS-CoV-2, Influenza, Monkeypox Virus, Zika virus, Dengue virus, and Poliovirus. Further investigation is needed into the co-infection’s role in SARS-CoV2-positive patients for a better understanding and management of the pandemic.

## Acknowledgments

We gratefully acknowledge the authors from the originating and submitting laboratories who generated and shared via GISAID genetic sequence data on which this research is based. We also thank Ms. Neha Kandpal, Ms. Jyoti, Mr. Nikil Gupta, Mr. Kaptan Verma, Mr. Namonarayan Meena for providing technical assistance and help in NGS library preparation and sequencing.

## Funding

Support from the Ministry of Health and Family Welfare, India.

## Author Contribution

Conceived of the study: SK, PR, HG, PM. Designed the study and experiments: KP, RVS, RM, MD, SK, PR Performed experiments: RVS, RM, MD, US, SK, KP. Collected and analyzed patient data: KP, RVS, RM, MD. Performed bioinformatics analyses: KP. Performed statistical analyses: KP, RVS, RM. Interpreted data: KP, RVS, RM, SK, PR.

## Conflict of Interest

The authors declare that they have no competing interests.

## Notes

### Competing Interest Statement

The authors have declared no competing interest.

